# Insulin-Like Peptides Regulate Feeding Preference and Metabolism in *Drosophila*

**DOI:** 10.1101/222539

**Authors:** Uliana Semanyuk, Dmytro Gospodaryov, Hrystyna Feden’ko, Ihor Yurkevych, Alexander Vaiserman, Kenneth Storey, Stephen Simpson, Oleh Lushchak

**Affiliations:** Department of Biochemistry and Biotechnology, Vasyl Stefanyk Precarpathian National University, 57 Shevchenka str., Ivano-Frankivsk, 76018, Ukraine; D.F. Chebotarev Institute of Gerontology, NAMS, 67 Vyshgorodska str., Kyiv, 04114, Ukraine; Institute of Biochemistry, Carleton University, 1125 Colonel By Drive, Ottawa, Ontario K1S 5B6, Canada; Charles Perkins Centre, The University of Sydney, NSW, 2006, Sydney, Australia

**Author notes:** These authors contributed equally to this article (Uliana Semanyuk, Dmytro Gospodaryov). Correspondence: Department of Biochemistry and Biotechnology, Vasyl Stefanyk Precarpathian National University, Ivano-Frankivsk, Ukraine. E-mail address (O. Lushchak).

**Keywords:** geometric framework, dietary response surface, macronutrient balance, nutrient intake trajectories, capillary feeding, Drosophila insulin-like peptides

## Abstract

Fruit flies have eight identified *Drosophila* insulin-like peptides (DILPs) involved in regulation of carbohydrate concentrations in hemolymph as well as accumulation of storage metabolites. In the present study, we investigated diet-dependent roles of DILPs encoded by genes *dilp1–5*, and *dilp7* in regulation of insect appetite, food choice, accumulation of triglycerides, glycogen, glucose, and trehalose in fruit fly body and carbohydrates in hemolymph. We found that *dilp2* gene predominantly influences body glycogen level, *dilp3* – trehalose level in hemolymph, while *dilp5* and *dilp7* affect triglyceride level. Fruit fly appetite was found to be regulated by *dilp3* and *dilp7* genes. Our data contribute to the understanding of *Drosophila* as a model for further studies of metabolic diseases and may serve as a guide for uncovering the evolution of metabolic regulatory pathways.

**HIGHLIGHTS:** Different Drosophila insulin-like peptides play distinctive roles in metabolism, physiology and appetite regulation.

Lack of Dilp2 and Dilp5 abrogates glycogen accumulation on high carbohydrate diets

Lack of Dilp3 leads to build-up of trehalose in haemolymph on high-carbohydrate-low-protein diets

Lack of Dilp3 and Dilp7 leads to increased consumption of protein on low-carbohydrate-high-protein diets

**GRAPHICAL ABSTRACT:** 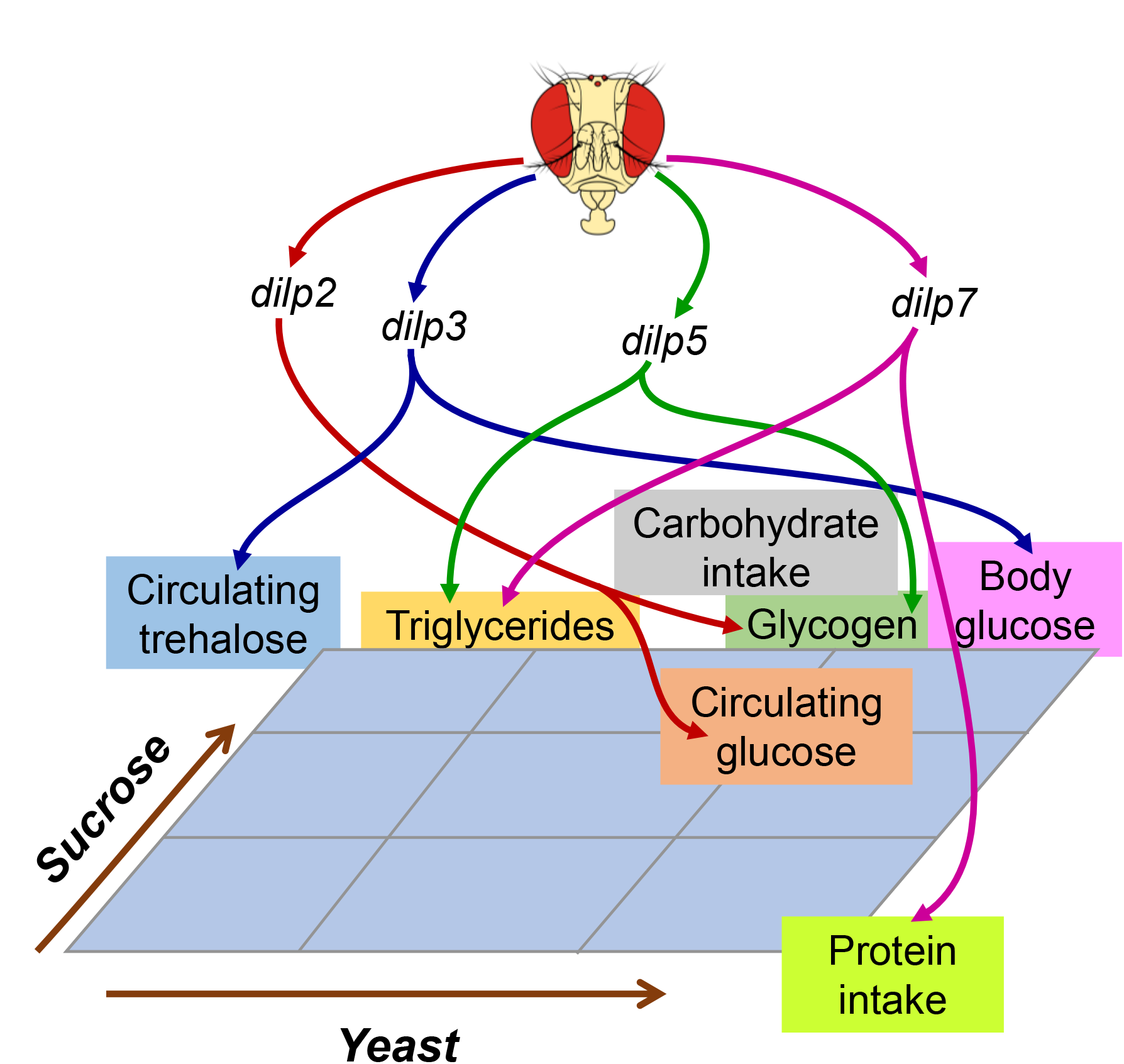

## INTRODUCTION

Fruit fly, *Drosophila melanogaster*, serves as a model for investigating the molecular mechanisms underlying pathological states such as type II diabetes, obesity, and metabolic syndrome. These conditions involve dysregulation of metabolism and share evolutionarily conserved genetic and environmental determinants (Leulier et al., 2017). Fruit flies and humans have common molecular regulators of key metabolic pathways, including hormones, transcriptional factors and signalling molecules which govern cellular and organ metabolism. They keep homeostasis and enable survival at a changeable environment, including diet.

Insulin plays a special role in metabolic regulation. This hormone regulates many points in carbohydrate and lipid metabolism, and disorders in its signalling and regulation cause diabetes. *Drosophila* insulin-like peptides (DILPs) regulate the balance between stored and circulating carbohydrates and also regulate metabolism by triggering signalling cascades that affect transcription of genes responsible for metabolic reconfigurations. Generally, DILPs promote accumulation of glycogen and fat stores (DiAngelo and Birnbaum, 2009), suppress gluconeogenesis but induce glycolysis in cells (Palanker Musselman et al., 2011). DILPs control growth and suppress food intake homeostatically when sufficient amount of food is ingested (Nässel et al., 2013). The disordering of such regulation leads to multiple defects of growth and development. Many recent investigations have revealed complex interactions between insulin signalling and other hormonal signals in *Drosophila*. Particularly, relationships have been shown between DILPs and developmental hormones of insects (Géminard et al., 2006), as well as small peptides which regulate appetite (Nässel et al., 2013). It is known that the release of DILPs is, in turn, controlled by neuropeptides with reverse feedback regulation (Nässel et al., 2013).

Unlike human insulin signalling, *D. melanogaster* has eight DILPs which are produced at different conditions and tissues, having diverse roles (Nässel et al., 2013). However, flies possess only one receptor which responds to all types of DILPs. Knockdowns of genes coding DILPs in *D. melanogaster* were shown to result in various growth defects (Zhang et al., 2009). Removal of insulin-producing cells in fruit flies phenocopy type I diabetes in humans (Rulifson et al., 2002). However, this has not yet been studied under different dietary conditions, where diet might modulate phenotype of DILP deficiency. Macronutrient combination, as well as the quality of macronutrients and interactions with micronutrients, can modulate disease outcomes, thus offering the opportunity for diet to serve as both prevention and cure (Simpson et al., 2015, Raubenheimer and Simpson, 2016, Simpson et al., 2017). We have shown recently in flies that macronutrient balance defines physiological parameters such as fecundity and stress resistance (Lushchak et al., 2014). It may alter also the impact of plant medicinal preparations (Gospodaryov et al., 2013). The goal of current study was to systematically vary dietary macronutrient balance in combination with specific DILP knockdowns to assess the effects on appetite and nutrient preferences of flies as well as on stored and circulating carbohydrates. This has allowed us to begin to dissect the roles of individual DILP types and to explore the dietary conditions under which different DILPs are released and operate.

## RESULTS

### DILPs Affect Protein Rather Than Sucrose Consumption

Individuals of the wild type line, *w^1118^*, and the mutant lines consumed larger volumes of solutions with low than high sucrose concentrations, indicative of compensatory feeding for sugar. Hence, wild type flies showed a decrease in consumption of sucrose solution when the concentration of carbohydrate was increased from 3 to 12% (*F_1,221_* = 438, *p* < 0.001; Fig 1A). The mutant lines all showed a similar trend, consuming lower volumes at higher carbohydrate concentrations. However *dilp5* and *dilp7* mutants ingested 25 and 30% more sucrose on 12S–3Y diet than the wild type, respectively. Flies of *dilp2* mutant line ate 14% more sucrose solution when given 12% solutions. Interactions on diet-dependent consumption of sucrose solution were observed between *dilp1* and *dilp7*, and *dilp3* and *dilp7* mutants (Table S1).

**Figure 1.**
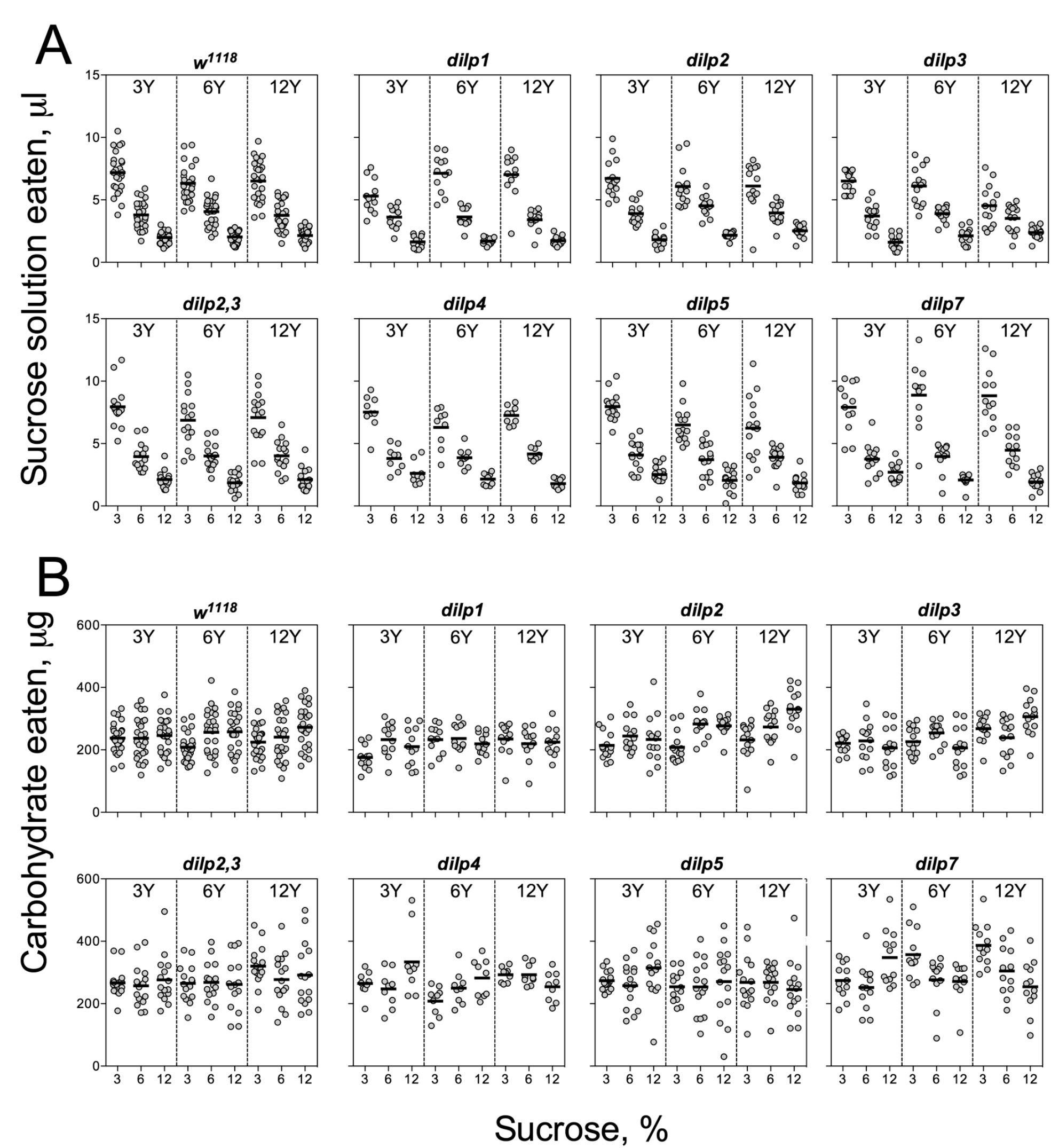
Consumption of carbohydrates by control and DILP-deficient fruit flies (A) Volumes of sucrose solution ingested. (B) Amounts of carbohydrate consumed. Tubes with tested fruit flies were supplied by pair of capillaries: one filled with solution of sucrose, and the other filled with solution of yeast autolysate. It was allowed flies to choose between capillaries for eating either yeast autolysate (mainly, source proteins and essential micronutrients such as vitamins, nitrogen bases, etc.) or sucrose (a source of carbohydrates). The combinations of 3%, 6%, and 12% sucrose, and 3%, 6%, and 12% of yeast autolysate were used. About 25- flies were tested condition.

The wild type flies demonstrated evidence for compensatory feeding for protein, although less marked than for carbohydrate, and consumed the biggest volume of yeast autolysate solution (Y) at its low concentrations (3% and 6%) when these were paired with a low concentration of sucrose (3%) (Fig. 2A). Mutations on *dilp1* and *dilp2* abrogated increased consumption of Y on 3S–3Y diet. Also, *dilp2* mutants consumed 28% less volume than the wild type on 6S–6Y diet. The wild type flies consumed 1.5-2.2-fold more yeast solution on diets with 3% sucrose than with 12% sucrose. This difference for *dilp2* mutants was 1.4-1.7-fold. The mutants on *dilp2* also consumed 1.4-2.1-fold more of Y than the wild type on diets with 12% Y. Flies of *dilp2,3* and *dilp7* lines ate 1.3-4.3-fold more Y on all diets as compared to the wild type flies. Mutants on *dilp4* and *dilp5* ingested 1.2-2.7-fold more Y than the wild type on all diets, except for 3S–6Y and 6S–6Y ones – for *dilp4* mutant, and 6S–3Y – for *dilp5* mutant.

**Figure 2.**
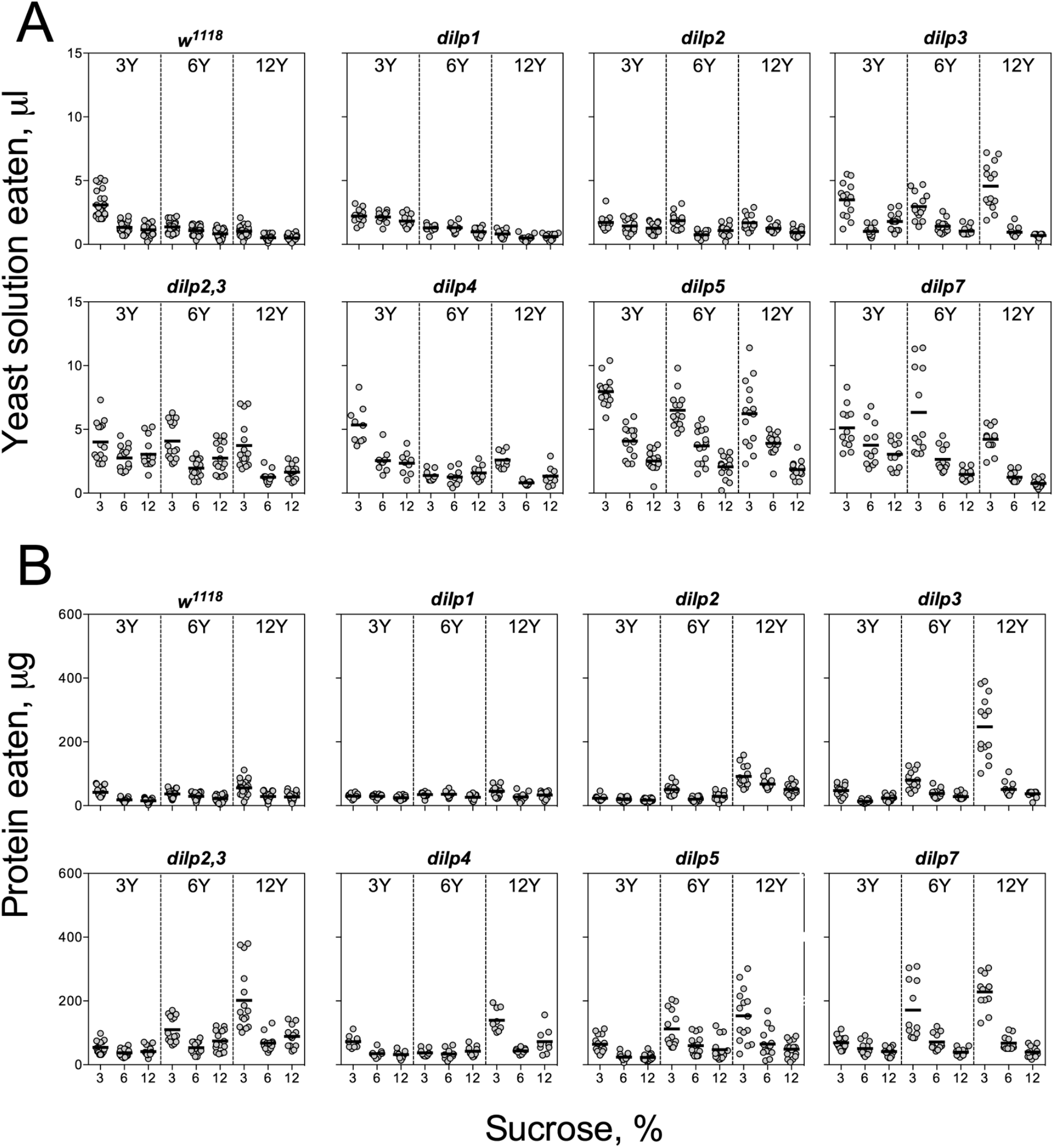
Consumption of protein-rich food source by control and DILP-deficient fruit flies (A) Volumes of yeast autolysate solution ingested. (B) Amounts of protein consumed.

Despite clear evidence for compensatory feeding for carbohydrate (above), the amount of consumed carbohydrate was not maintained constant across food treatments and was dependent on sucrose concentration for the wild type flies (*F_1,221_* = 11.5, *p* < 0.001), being higher on the high as compared to the low sucrose solutions (Fig. 1B). Dependence of the amount of consumed carbohydrate on sucrose concentration was also seen for *dilp2, dilp3, dilp4, dilp5*, and *dilp7* lines (Table S2). However, carbohydrate consumed by the mutant on *dilp2,3* was dependent on the concentration of yeast provided (*F_1,131_* = 4.72, *p* < 0.05). The mutants on *dilp2,3, dilp4*, and *dilp7* ingested 21%, 26% and 58% more carbohydrate on diet 3S–12Y than the *w^1118^* flies, respectively. Flies of *dilp2,3, dilp5* and *dilp7* mutant lines consumed 1.3-, 1.2-, and 1.6-fold more carbohydrates than the wild type on 3S–6Y diet. The mutations on *dilp4, dilp5*, and *dilp7* increased carbohydrate consumption by 25-33% on 12S–3Y diet. The *dilp4* and *dilp7* mutants ate 13% and 19% more carbohydrates on 6S–12Y diet as compared to *w^1118^* flies, respectively. Somewhat higher carbohydrate consumption (15% and 21%) was observed for *dilp5* flies on 3S–3Y and 3S–6Y diets.

The pattern for amount of protein consumed followed the concentration of Y provided. The *dilp2,3* and *dilp7* mutants consumed 1.3-4.3-fold more protein on all diets as compared to the wild type. Mutants on *dilp2, dilp2,3, dilp3, dilp4, dilp5*, and *dilp7* ingested 1.4-, 3.0-, 4.3-, 2.3-, 2.7-, and 4.3-fold greater amount of protein on 3S–12Y diet than the wild type flies, respectively (Fig. 2B). Similarly, *dilp2, dilp3, dilp2,3, dilp5*, and *dilp7* mutants demonstrated 1.3-, 2.0-, 2.5-, 2.1-, and 3.3-fold higher amounts of ingested protein on 3S–6Y diet as compared to the wild type.

Knowing the volumes of the different solutions consumed and the concentrations of nutrient within the solutions, we were able to plot the relationship between protein and carbohydrate eaten using nutritional geometry (Fig. 3). Small differences between lines were observed on 3S–3Y and 12S–3Y diets. However, flies tended to eat more carbohydrate than protein, keeping the P:C ratio of ingested nutrients as 1:8 or 1:4. On the 3S–3Y diet, *dilp2* mutant kept the P:C ratio of 1:8, while *dilp1, dilp4*, and *dilp7* mutants kept 1:4 ratio. The other lines, including wild type, kept P:C ratio between 1:8 and 1:4. On 12S–3Y diet, trajectories for all lines converged close to a 1:8 ratio. On 3S–12Y diet, wild type and *dilp1* mutant converged on P:C ratio 1:4. On this diet, P:C trajectories for *dilp2, dilp2,3, dilp4, dilp5* and *dilp7* mutants were skewed towards a ratio of 1:2, while *dilp3* mutant flies ingested even more protein and less carbohydrate, reaching a 1:1 ratio. Of note, flies of *dilp7* line consumed the largest amount of carbohydrate among all lines on 3S–12Y diet and also rather large amounts of protein. Increase in sugar concentration on 12S–12Y diet leveled the effect observed for mutant lines on 3S–12Y diet. These observations indicate that when wild type flies restrict sipping from autolysate-containing capillary, the mutant lines sipped equally from both capillaries.

**Figure 3.**
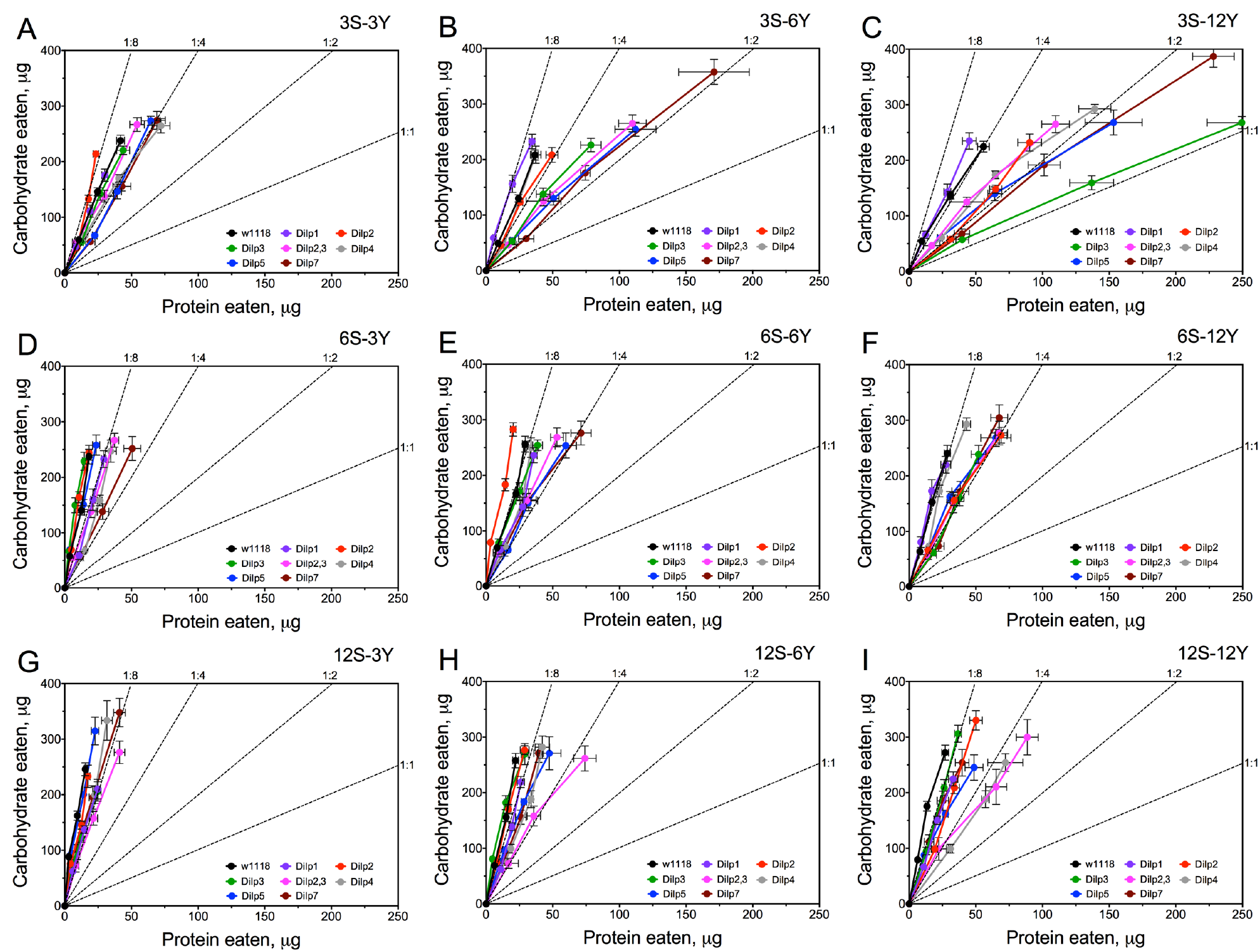
Nutrient intake trajectories resulting from volumes ingested of the two food solutions (sucrose and yeast) for all fruit fly lines tested

### Different DILPs Influence Hemolymph Glucose in a Diet-dependent Manner

Hemolymph glucose (HG) in wild type flies was affected by diet and was dependent on dietary sucrose (*p* = 0.026) rather than yeast (*p* = 0.601) or their interaction (*p* = 0.130) (Fig. 4A; Table S3). Lack of Dilps reshapes the relationship between HG and dietary components. Particularly, in *dilp3* and *dilp7* mutant flies HG was dependent on dietary yeast (*p* = 0.001 and 0.012 respectively). Neither, yeast nor sucrose affected HG in *dilp2* and *dilp5* mutant flies.

**Figure 4.**
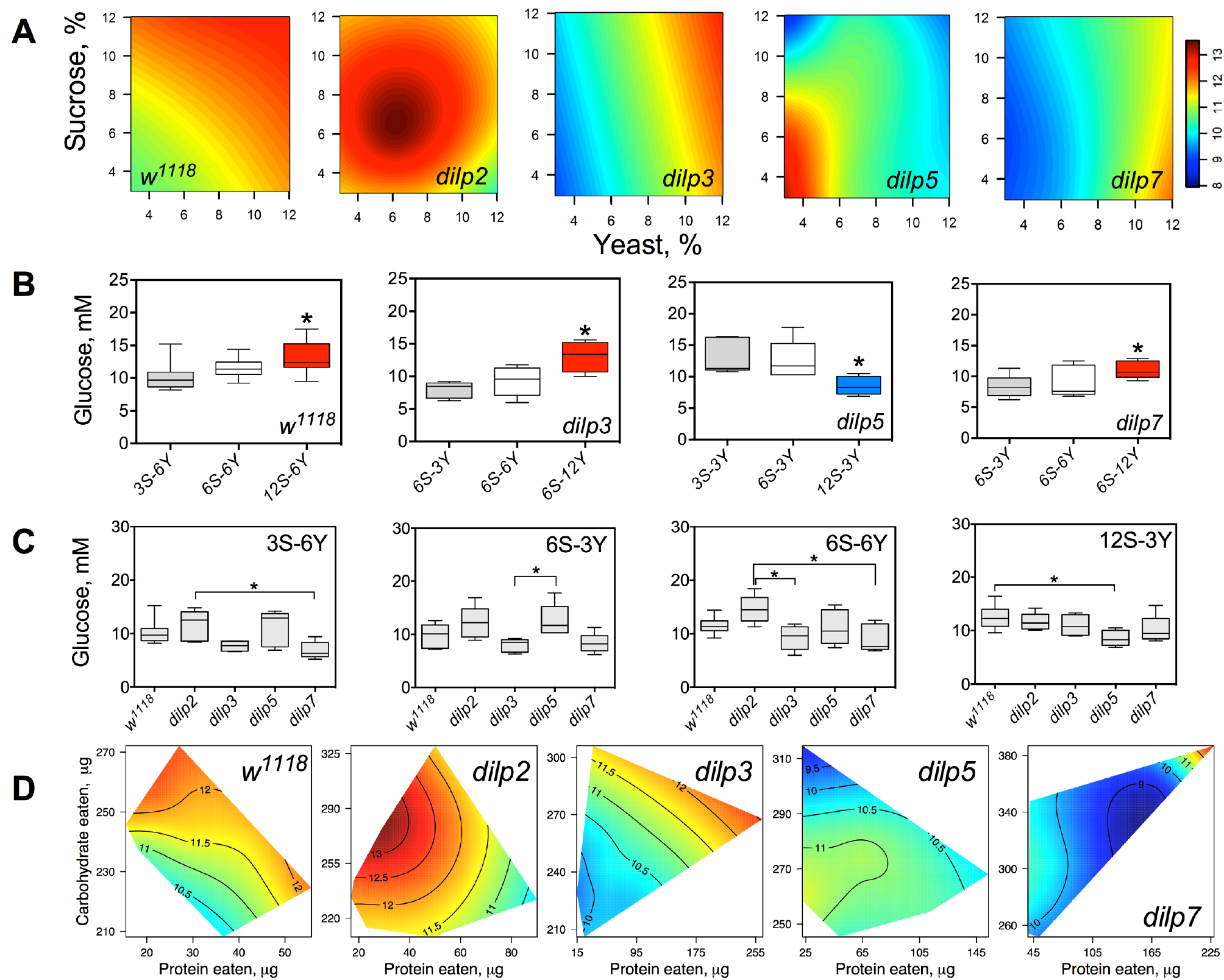
dlLPs deficiency changes the diet-dependent pattern of glucose concentration in fruit fly hemolymph (A) Dietary response surfaces depicting dependence of concentration of glucose in fly hemolymph (mM) on concentration of yeast and sucrose in the diet. There were tested nine combinations of sucrose (3%, 6%, and 12%) and yeast (3%, 6%, and 12%). In each case, the flies of tested lines were able to choose between of sucrose- or yeast-containing medium. All five surfaces are placed under one scale. (B) The remarkable differences between hemolymph glucose values in flies of each line kept on different diets. (C) The remarkable differences between hemolymph glucose values in flies of different lines on particular diets. (D) Dietary responses surface depicting dependence of hemolymph glucose in wild type and *dilp* mutant lines on amounts of protein and carbohydrate consumed.

The wild type flies had by 27% higher HG on 12S-6Y diet than those fed 3S-6Y diet (Fig. 4B). HG was 57% and 31% higher on the diet 6S-12Y compared with 6S-3Y in *dilp3* and *dilp7* mutants, respectively. *dilp5* mutant flies had lower HG by 36% when fed 12S-3Y diet compared to 3S-3Y. The differences in HG in flies of different genotypes were observed on distinct diets (Fig. 4C). Particularly, *dilp7* mutant flies had 14% and 20% lower HG on diets 3S-6Y and 6S-6Y in comparison with values for *dilp2* mutant flies. In addition, HG was lower by 35% in *dilp3* mutants compared to *dilp2* on 6S-6Y diet. The *dilp5* mutants had 32% higher HG than *dilp3* flies fed 6S-3Y diet. In addition, loss of *dilp5* significantly decreased HG on high-sucrose low-yeast diet 12S-3Y.

Loss of DILPs affected the HG in response to protein and carbohydrate intake (Fig. 4D). The highest HG in wild type flies was observed at high-carbohydrate and low protein intake. In *dilp2* mutant flies, HG was maximized at consumption of moderate amounts of sucrose and low amounts of protein. Maximum values for HG were observed in *dilp3* mutants that consumed high amounts of protein and moderate amounts of carbohydrate. Oppositely, the highest HG in *dilp5* mutants was observed at consumption of moderate amounts of carbohydrate and was independent of amount of protein consumed. Mutation on *dilp7* resulted in highest HG at high protein and high carbohydrate intakes.

### Dilp3 Regulates Hemolymph Trehalose on Low-Protein and High-Carbohydrate Diet

Concentration of trehalose in hemolymph (HT) of wild type flies was dependent on the concentration of dietary yeast (*p* = 0.001) and was modulated by sucrose (*p* = 0.013 for the interaction term between yeast and sucrose; Fig. 5A; Table S4), although sucrose concentration as a main effect did not influence HT (*p* = 0.557). HT in *dilp2* mutant was dependent on sucrose (*p* = 0.002). Whereas there was no main effect of yeast (*p* = 0.9517), there was a statistically significant interaction between sucrose and yeast concentration (*p* = 0.005). Concentration of hemolymph trehalose in *dilp3* mutants was strongly dependent on dietary yeast (*p* = 0.001) and sucrose (*p* < 0.001). A weak dependence of HT on dietary sucrose was shown for the *dilp5* mutant line (*p* = 0.049). Indeed, the HT values of *dilp5* mutants were very similar in flies kept on all diets. There was also dependence of HT in *dilp5* mutants on both nutrients (*p* = 0.032). A strong dependence on concentration of dietary yeast was found for the *dilp7* mutant (*p* < 0.001).

**Figure 5.**
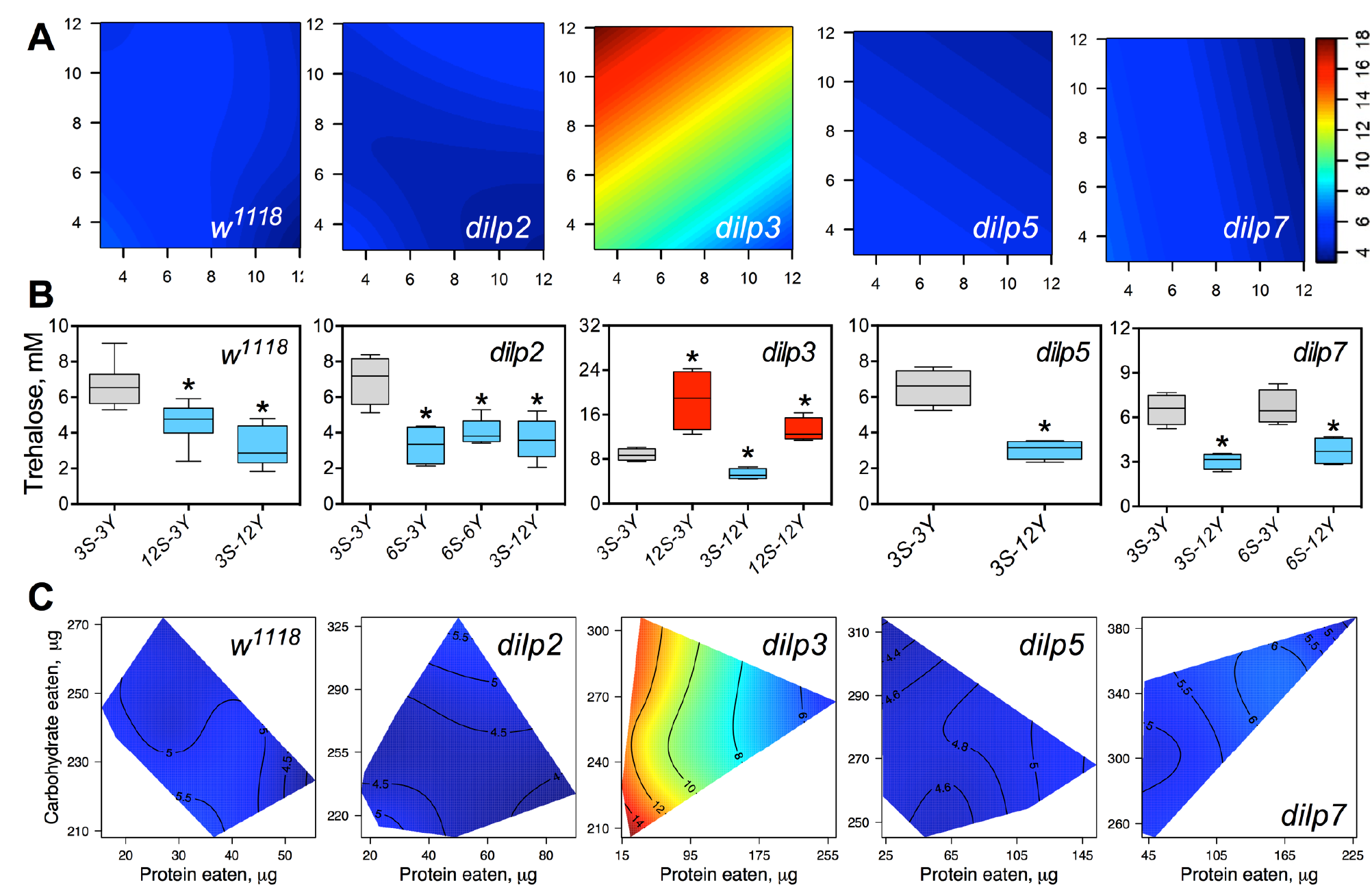
The effect of mutations on *dilp* genes on the diet-dependent pattern of trehalose concentration in fruit fly hemolymph (A) Dietary response surfaces representing dependence of concentration of trehalose in fly hemolymph (mM) on concentrations of yeast and sucrose in the diet. There were tested nine combinations of sucrose (3%, 6%, and 12%) and yeast (3%, 6%, and 12%). In each case, the flies of tested lines were able to choose between of sucrose- or yeast-containing medium. All five surfaces are placed under one scale. (B) The remarkable differences between hemolymph trehalose values in flies of each line kept on different diets. (C) Dietary response surfaces showing dependence of hemolymph glucose in wild type and *dilp* mutant lines on amounts of protein and carbohydrate consumed. Each surface has own scale shown by contour lines.

The wild type flies fed 3S-12Y diet had 2.3-fold lower values than those kept on 3S–3Y diet (Fig. 5B). We observed 27% and 32% lower HT values in the wild type flies fed on 12S-3Y and 12S-12Y diets, respectively, when compared to 3S-3Y diet (Fig. 5B). The *dilp2* flies kept on 3S–3Y diet had 2.1-, 1.9-, and 2.0-fold higher concentration of hemolymph trehalose than flies fed on 6S–3Y, 6S–6Y, and 3S–12Y diets, respectively. In *dilp3* line, the lowest HT value was observed on 3S-12Y diet, being 42% lower compared to that on 3S-3Y diet. The *dilp3* flies fed 12S-3Y diet had 2.2-fold higher trehalose concentration in hemolymph than those fed 3S–3Y diet, while those fed 12S-12Y diet showed 1.4- and 2.5-fold higher HT than the flies on 3S–3Y and 3S–12Y diets, respectively. Only those *dilp5* flies that consumed 12S-3Y diet had lower (29%) hemolymph trehalose than flies on 3S-3Y diet. The *dilp7* flies on 3S-12Y and 6S-12Y diets contained 2.1- and 1.7-fold less trehalose in hemolymph than, respectively, on 3S-3Y and 6S–3Y diets.

Interestingly, *dilp3* flies had 1.6-4.0-fold higher HT values compared to the wild type line (Fig. 5A, 5C). The smallest difference between the lines was observed on 6S-6Y diet, while the largest differences occurred on 12S–3Y diet. Trehalose concentration in hemolymph was not significantly different between *w^1118^* and *dilp5*, and *dilp7* flies, while flies of *dilp2* line had 41% lower hemolymph trehalose concentration on 6S–3Y diet than *w^1118^* flies.

A clear dependence of hemolymph trehalose concentration on amounts of macronutrients eaten was seen only for *dilp3* and *dilp7* mutants (Fig. 5C). Particularly, *dilp3* mutant flies had the highest concentration of trehalose in hemolymph only when consuming low amounts of protein. The concentration was only marginally influenced by amount of carbohydrate consumed. The opposite situation was observed for *dilp7* mutant. The peak in concentration of circulating trehalose was observed at high amounts of carbohydrate and protein consumed. At the same time, the lowest values of circulating trehalose were observed at low amounts of protein and carbohydrate eaten.

### Dilp2 and Dilp5 Regulate Glycogen Accumulation Independently of Diet

Glycogen content was dependent on sucrose concentration in wild type flies (*p* = 0.002; Fig. 6A; Table S5) and in *dilp3* and *dilp7* flies. Glycogen content in the *dilp7* mutant resembled the diet-dependent pattern attributed to the mutant on *dilp3*. Mutations on *dilp5* and *dilp7* made glycogen accumulation also dependent on both yeast and sucrose (*p* = 0.031 and *p* = 0.004 respectively), with also a strong effect of dietary yeast concentration for *dilp5* mutant (*p* = 0.027) (Table S5). However, glycogen content of *dilp2* mutant was dependent on neither dietary sucrose, nor yeast, or their combination.

**Figure 6.**
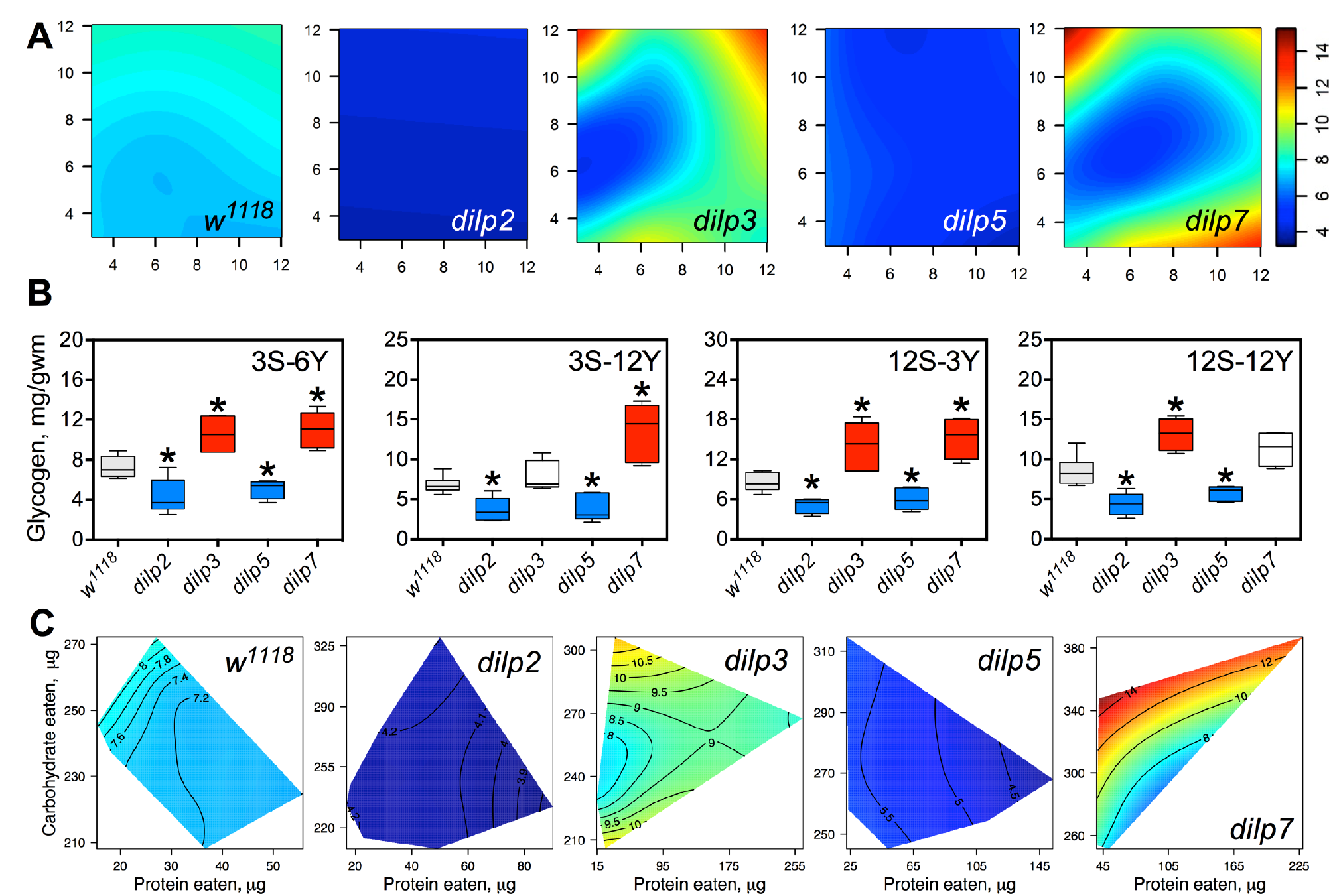
The effect of *dilp* mutations on the diet-dependent pattern of glycogen accumulation in fruit fly body (A) Dietary response surfaces depicting dependence of glycogen content in the body of individuals of wild type (*w^1118^*) and *dilp* mutant lines on concentrations of yeast and sucrose in the diet. (B) The remarkable differences in glycogen content between investigated fruit fly lines. (C) Dietary response surfaces showing dependence of glycogen content in wild type and *dilp* mutant lines on amounts of protein and carbohydrate consumed. Each surface has own scale shown by contour lines.

The mutants on *dilp2* had 1.5- to 2.4-fold lower glycogen content on all diets than the wild type line (Fig. 6A). The smallest difference between these lines was observed on 12S–3Y diet, while the largest difference was on the 12S-6Y diet (Fig. 6B). The mutant on *dilp3* accumulated larger amounts of glycogen on diets 3S-6Y, 12S-3Y and 12S-12Y than the wild type. The *dilp5* mutant had 1.3- to 2.2-fold lower glycogen content than the wild type on high sugar (12S) and high yeast diets (12Y) (Fig. 6B). The biggest difference between these lines was observed on the diet 3S-12Y. The *dilp7* mutant accumulated higher than the wild type amounts of glycogen on the diets 3S-6Y, 3S-12Y, and 12S-3Y.

Significantly, 1.9-fold higher glycogen content was observed on the 12S-12Y diet compared to 3S-12Y in *dilp3* flies, while *dilp7* flies accumulated by 2.0-fold more glycogen on 12S-3Y diet compared to 3S-3Y diet.

Amount of stored glycogen increased with the increase in amount of consumed carbohydrate in wild type flies (Fig. 6C). Deficiency in particular Dilps changed this pattern. In *dilp3* mutants, dependence of glycogen accumulation on amount of carbohydrate consumed was disruptive, being maximal at either very low or very high amounts of eaten carbohydrates. In *dilp5* mutants, amount of glycogen decreased with the amount of protein eaten but did not depend on amount of carbohydrate eaten. Consumption of carbohydrates promoted glycogen synthesis in *dilp7* mutants. However, protein consumption led to lowering of glycogen accumulation.

### Dilp3, Dilp5 and Dilp7 Affect Accumulation of Triglycerides

The wild type flies and *dilp2* mutant flies accumulated high amounts of triglycerides (TAG) on high carbohydrate diets, strongly dependent on dietary sucrose (*p* = 0.0003 and *p* = 0.0001, respectively; Fig. 7A; Table S6). Content of TAG in *dilp2* mutant was dependent also on dietary yeast (*p* = 0.0123). A similar effect was characteristic for *dilp7* flies, in which TAG content was also affected by both nutrients ((*p* = 0.0194 and 0.0168 for sucrose and yeast, respectively).

**Figure 7.**
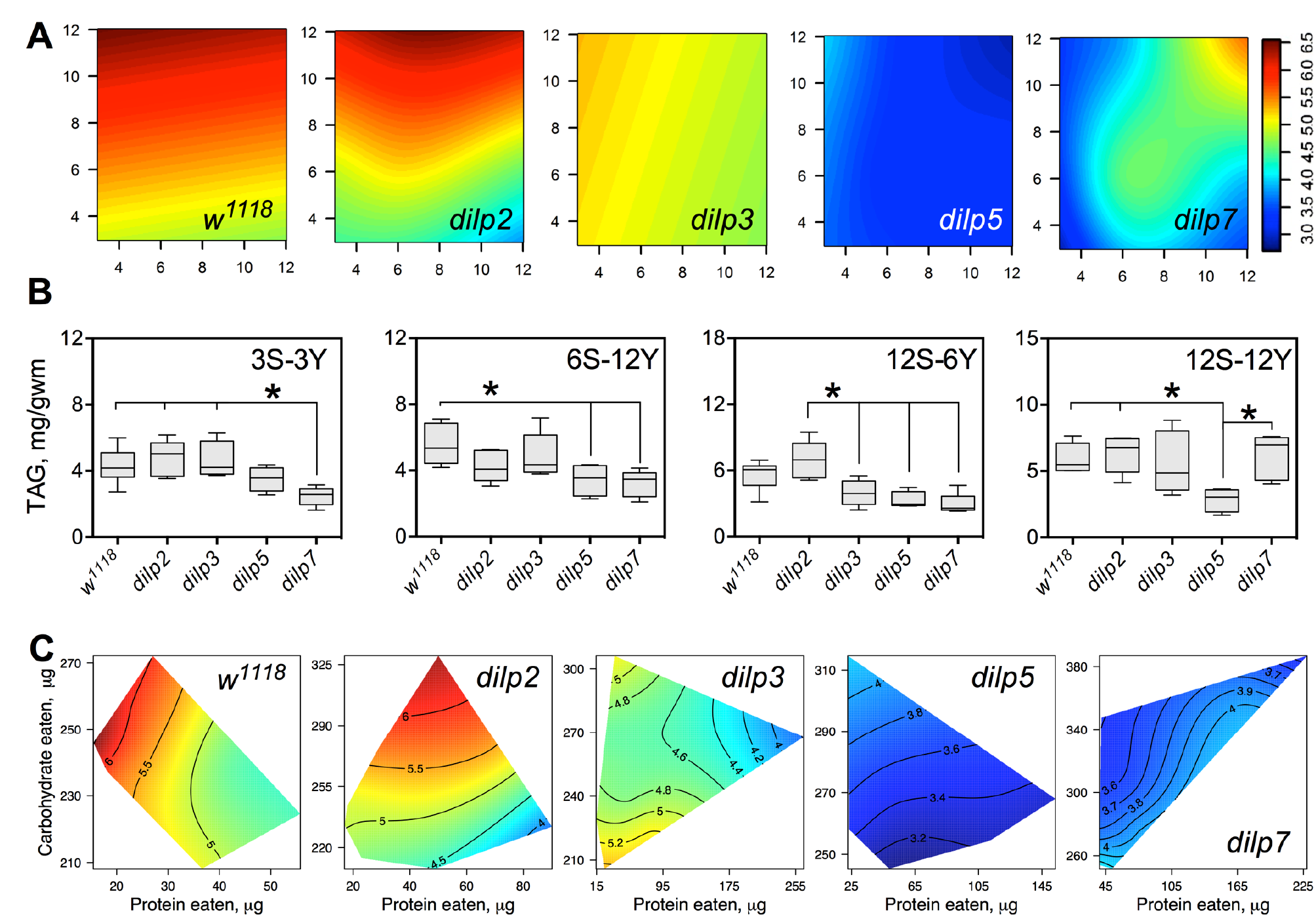
The effect of *dilp* mutations on the diet-dependent pattern of triacylglyceride (TAG) accumulation in fruit fly body (A) Dietary response surfaces depicting dependence of TAG content in the body of individuals of wild type (*w^1118^*) and *dilp* mutant lines on concentrations of yeast and sucrose in the diet. (B) The remarkable differences in TAG content between investigated fruit fly lines. (C) Dietary response surfaces showing dependence of TAG content in wild type and *dilp* mutant lines on amounts of protein and carbohydrate consumed. Each surface has own scale shown by contour lines.

The wild type flies had 74% more TAG on 12S-3Y compared to 3S-3Y diet. In addition, these flies had 21% more TAG on 12S-12Y diet compared to 3S-12Y. The difference was less pronounced on diets 3S-6Y and 12S-6Y. The *dilp2*-deficient flies showed similar to the wild type pattern of diet-dependent TAG accumulation. However, difference in TAG content between low and high sugar diets was more marked than in the wild type, having been shifted more to the high yeast diets. It was 2.4-fold between 3S-12Y and 12S-12Y diets, and 1.7-fold between 3S-6Y and 12S-6Y diets.

Mutations on *dilp3, dilp5* and *dilp7* changed the pattern of TAG accumulation on high sugar diets. Flies of *dilp5* line had 1.5-2.3-fold lower amount of TAG than the wild type on all high sucrose diets, and on diets 6S-6Y, 6S-12Y, and 3S-6Y (Fig. 7B). The *dilp3* mutants showed 1.6-and 1.7-fold lower amounts of TAG than the wild type and *dilp2* mutant on 12S-6Y, respectively. The *dilp7* mutant had 1.5-1.9-fold lower TAG content than *w^1118^* flies on all low yeast diets. A 1.4-, 1.5-, and 2.4- lower TAG content compared to the wild type was observed on 3S-12Y, 6S-12Y and 12S-6Y diets, respectively.

In many cases, flies reared on diets with high carbohydrate content accumulated the highest amounts of TAG (Fig. 7C). When we assess dependence of TAG content on amounts of carbohydrate and protein eaten, we see clear decrease of TAG content in body at high amounts of protein consumed in wild type flies and *dilp2* mutants.

### Mutations on *dilp3* and *dilp7* Lead to Increased Body Glucose on High-Yeast Diets

The concentrations of dietary sucrose and yeast had a relatively weak influence on the concentration of glucose in fly body, and this was observed for all lines. A dependence on sucrose concentration was exhibited by the wild type and *dilp7* lines (*p* = 0.041 and 0.0064, respectively; Fig. S1A, Table S7). A dependence of body glucose on yeast concentration was observed for the *dilp3 (p* = 0.018) and *dilp7* mutants (*p* = 0.010).

The wild type flies had 34% higher level of body glucose on 12S-12Y diet as compared to 12S-3Y (Fig. S1B). Similarly, *dilp7* flies accumulated 41% more glucose on 12S-12Y diet than on 12S-3Y diet. The *dilp3* mutant flies showed significant difference on the parameter only between 3S–3Y and 3S–6Y diets. The *dilp2* and *dilp5* mutant flies had 25% and 21% lower level of glucose in bodies on 3S–6Y diet versus wild type flies, respectively. Interestingly, *dilp3* and *dilp7* mutant flies had 1.3- and 1.8-fold higher levels of body glucose on all diets with 12% sucrose and on 3S-6Y and 3S-12Y diets compared to the wild type flies. Additionally, *dilp3* mutants had a 1.6-fold higher level of body glucose on 6S-3Y diet.

The concentration of glucose in the fly body reached the highest value on high intakes of protein in wild type line and was dependent on amount of consumed protein in general (Fig. S1C). A somewhat similar situation was observed for *dilp3* line. However, in *dilp3* flies the transition from low to high values of body glucose was steeper than in *w^1118^* flies. The highest values were observed at high levels of both consumed protein and carbohydrate. A similar pattern was also observed in *dilp7* mutant, however the lowest value was observed at moderate intakes of protein, unlike *w^1118^* and *dilp3* flies where the lowest value of body glucose corresponded the lowest intakes of protein.

### All *dilp* Mutations Lead to Specific Body Trehalose Response to Diet

Body trehalose was slightly affected by diet in wild type flies. The lowest value was observed on 6S-12Y diet. The highest amount of body trehalose was observed on 3S-12Y and 12S-3Y diets. Mutation on *dilp2* leveled these diet responses observed in wild type flies, leading to virtually equal accumulation of trehalose in fruit fly body independently of diet (Table S8). Mutation on *dilp3* led to a relatively small decrease in levels of body trehalose on diets with low concentrations of yeast extract. However, mutation on *dilp5* caused a drastic decrease in body trehalose levels on diets with either low (3%) or high (12%) concentrations of sucrose (Fig. S2). Mutants on *dilp7* exhibited similar patterns of body trehalose to wild type line but with more pronounced increase of body trehalose on 3S-3Y, 3S-6Y, and 3S-12Y diets.

### Mutations on DILPs Did Not Affect the Influence of Diet on Body Mass

Body mass was dependent on dietary sucrose in wild type flies (*p* = 0.022) with a marginally statistically significant effect of dietary yeast (*p* = 0.049) (Table S9). There were no significant effects of diet on body mass in *dilp* mutants.

## DISCUSSION

### Food Consumption and Appetite

As in previous studies, we have found that fruit flies consume more food on low than on higher percent carbohydrate diets, indicative of compensatory feeding for carbohydrates (Lushchak et al., 2011; Lushchak et al., 2012; Lushchak et al., 2014; Lushchak et al., 2016, Rovenko et al., 2015a). DILPs are known to be connected with signaling pathways that control appetite (Wu et al., 2005; Itskov and Ribeiro, 2013; Nässel et al., 2013; Luo et al., 2014; Lushchak et al., 2015), and diminution of insulin signaling (IS) leads to increase in consumption of unpalatable food by fruit flies or their larvae (Wu et al., 2005; Luo et al., 2014). Lack of functional Dilp4, Dilp5 and Dilp7 led to significantly increased food consumption on almost all diets. Lack of either Dilp2 or Dilp3 led to increase in food consumption only on high protein diets. However mutation in both genes resulted in increased food consumption on all diets, implying that the actions of Dilp2 and Dilp3 are somewhat synergistic. Notably, *dilp7* mutants had the most pronounced increase in food intake on low sucrose concentrations, and also showed compensatory responses to changes in concentration of the yeast solution, for example consuming substantially more volume of food than wild type flies on the 3S-3Y and the 3S-6Y treatments. Thus, *dilp7* appears to be the most associated among the DILPs in regulation of macronutrient intake. Dilp7 is believed to be a relaxin-like peptide in *Drosophila*, regulating egg laying decisions. It is notable that protein and carbohydrate intake are essential for reproduction, with protein appetite being tightly coupled to mating and egg production (Lee et al. 2008; Sisodia and Singh, 2012; Simpson and Raubenheimer, 2012; Lushchak et al., 2014; Liu et al., 2017, Leitão-Gonçalves et al., 2017). Interestingly, human relaxins were shown to be orexigenic (McGowan et al., 2008; Grosse et al., 2014). Partial co-expression of *dilp7* with short neuropeptide F involved in feeding behavior (Nässel et al., 2013) was recently shown. Additionally, Dilp7 has been reported to be involved in regulation of feeding behavior by other authors (Owusu-Ansah and Perrimon, 2014). Increased appetite was characteristic also for other DILP mutants, though to a lesser extent than for Dilp7. This may suggest that other DILPs also function at least partially as relaxin-like peptides. However, it poses the question as to whether specific DILPs, especially Dilp7, may bind other receptors than *Drosophila* insulin receptor (dInR). Human relaxins and insulin-like peptides are known to bind specific G-protein coupled receptors distinct from tyrosine kinase type insulin receptor (Bathgate et al., 2013).

It was also found recently that responses on continuous lack of protein and carbohydrates are mediated by distinct dopaminergic neurons (Liu et al., 2017). Moreover, activation of protein feeding simultaneously inhibited carbohydrate feeding. Our results add complexity to these data, suggesting that the response is additionally mediated by DILPs.

### Glucose and Trehalose

The only DILP mutation which led to substantial increase in the level of hemolymph glucose was Dilp2. Lack of other DILPs resulted in a decrease in the level of hemolymph glucose on some diet treatments and an increase on others (e.g., *dilp5* mutant). The effect suggests that Dilp2 may physiologically be the closest DILP to human insulin. The regulation of hemolymph glucose level by Dilp2 is also confirmed by other authors (Zhang et al., 2009), while its physiological proximity to insulin is supported by amino acid sequence comparison (Brogiolo et al., 2001), and activation of dInR phosphorylation by Dilp2 binding (Rulifson et al., 2002). The dependence of hemolymph glucose on protein intake in *dilp3, dilp5*, and *dilp7* mutants suggests that DILPs may regulate distinct branches of glucose metabolism. Particularly, the level of circulating glucose may depend on the balance between cellular influx and efflux of glucose, glycogen synthesis and breakdown, synthesis of glucose from organic acids via gluconeogenesis, and on glucose catabolism via glycolysis and/or pentose phosphate pathways. In fruit flies, glucose can also be used for trehalose synthesis. Re-routing of glucose metabolism is possible for *dilp5* and *dilp7* mutants since we did not see increase in the levels of other metabolites investigated on the treatments where these mutants had low levels of hemolymph glucose. However, it was seen that circulating trehalose levels showed an inverse relationship to circulating glucose in *dilp3* mutant (Fig. 3C versus Fig. 4D for *dilp3* mutant). This suggests that a lack of Dilp3 may result in conversion of glycogen stores into trehalose instead of glucose. This possible conversion was inhibited by increased intake of protein. Therefore, our current data strongly implicate Dilp3 in regulation of trehalose concentration in hemolymph. This finding is not consistent with previous data, which showed that knockdown of *dilp2* gene but not *dilp3* gene led to an increase of trehalose content in whole fly (Broughton et al., 2008; Kannan and Fridell, 2013), however, the diet used was more concentrated than most of the diets in the current study. Interestingly, *dilp3* mutants also had high level of glucose in the body. The latter may imply that Dilp3 would normally suppress gluconeogenesis and trehalose synthesis. In case of Dilp3 lack, consumed carbohydrates would be re-directed for trehalose synthesis. The similar pattern of diet-dependent accumulation of trehalose in the body was observed for *dilp7* mutant. In *dilp7* mutant, this accumulation was not, however, reflected by an increase in the level of hemolymph trehalose, suggesting that *dilp7* is not involved in regulation of this branch of metabolism.

### Glycogen and Triacylglycerides

Using nutritional geometry, we have shown that both *dilp2* and *dilp5* are involved in regulation of glycogen synthesis in fruit flies, a function that has not been reported previously for these DILPs (Broughton et al., 2008; Grönke et al., 2010). Glycogen along with proteins also contributes to fly weight, which is reflected in our study where *dilp2* mutants were significantly lighter than controls on high carbohydrate diet. Earlier, it was shown that *dilp5* expression is dependent on dietary carbohydrate (Morris et al., 2012; Pasco and Léopold, 2012). On the other hand, previously we did not find significant change in *dilp5* expression with an increase in sugar concentration in larval diet (Rovenko et al., 2015a). It was, however, dependent on the type of carbohydrate: increase in glucose concentration did not change *dilp5* expression, while fructose suppressed steady-state levels of *Dilp5* transcript (Rovenko et al., 2015b).

As we can see from the current data, TAG and glycogen accumulation were most pronounced on high sucrose diets in control flies. Lack of Dilp5 led to decrease in TAG content for all treatments. Hence, Dilp5 seems to be especially necessary for nutrient sensing on high carbohydrate and high yeast diets, and also for converting ingested food into carbohydrate and lipid storage on these diets. Recent data on *dilp5* expression confirm these suggestions. Particularly, expression of *dilp5* was dependent on both consumed carbohydrate and protein in the study of Post and Tatar (2016), while triggered by yeast in larvae in the study of Okamoto and Nishimura (2015).

The role of Dilp5 in regulation of glycogen and TAG synthesis/breakdown has not been reported previously. Our study design was notable in two respects. First, we allowed flies to choose between protein and carbohydrate sources. This choice takes into account an impact of a *dilp* mutation on nutrient sensing. Indeed, *dilp* expression and release of the peptides from neurosecretory cells is dependent on availability of particular nutrients. The sensory response to particular nutrients which is transduced to insulin-producing cells was shown to be mediated by biogenic amines (Luo et al., 2014) and food odors (Lushchak et al., 2015). Secondly, we created conditions where flies were relatively freed to fly and otherwise move around, thereby expending energy and creating a demand for glycogen and TAG catabolism.

As compared with control flies, lowered levels of TAG were also found in the *dilp7* mutants. The effects of *dilp7* knockout on lipid metabolism are indirectly confirmed by recent data, showing that down-regulation of insulin signaling led to reduced lipid accumulation (Linneweber et al., 2014). In previous studies, ablation of insulin-producing cells either resulted in no change in lipid content (Broughton et al., 2008) or slightly increased it (Broughton et al., 2005).

### General Picture and Conclusions

The present study explored the influence of *dilp* knockouts on 1) food intake and 2) levels of stored and circulating metabolites. The functions of DILPs in *Drosophila* resemble that of insulin in mammals, namely the lowering glucose level in blood (hemolymph in insects) by increasing glucose uptake by cells and directing it for production of storage metabolites, including glycogen and lipids. Our current data showed that the balance of dietary macronutrients, proteins and carbohydrates, influences the outcomes of an insulin-like peptide deficiency. Moreover, we have shown that knockouts on insulin-like peptides of *Drosophila* affect feeding behavior. It is clearly seen that different DILPs regulate particular aspects of diet-dependent metabolism. From the current data, it is seen that Dilp2 and Dilp5 can be involved in diet-independent accumulation of glycogen. Dilp3 was shown to be responsible for regulation of trehalose synthesis and/or release of trehalose into hemolymph, while Dilp5 and 7 were responsible for TAG synthesis on high carbohydrate diets. These specific roles for DILPs can be conferred by likely nutrient-dependent DILP release by neurosecretory cells and specific target cells for DILPs. The specific connection between feeding behavior and DILPs, found in the current study, may reflect an impact of nutrient cues on signaling effects of DILPs. In other words, specific ratios of macronutrients can evoke DILP release and subsequent regulation of metabolism in particular tissues.

## EXPERIMENTAL PROCEDURES

### Fruit Fly Lines

Flies *Drosophila melanogaster* were cultured on standard medium (7.5% molasses, 5% yeast, 6% corn, 1% agar and 0.18 nipagin as a mold inhibitor) at 25°C, 12:12 light:dark cycle and 60% relative humidity. Flies of the *w^1118^* line and mutants generated on the *w^1118^* background for *dilp1, dilp2, dilp2,3, dilp3, dilp4, dilp5* and *dilp7* were kindly provided by Dr. Sebastian Grönke (Max Planck Institute for Biology of Ageing, Cologne, Germany) (Grönke et al., 2010).

### Feeding Assays

For experiments, two-day-old flies were transferred to fresh food and kept for an additional 48 h for mating. Females were collected under light CO_2_ anesthesia and placed into plastic tubes (1.5 ml) with three holes for ventilation, and two holes in the cap for the positioning of 5 μl capillary feeding tubes (Drummond Scientific Company) filled with nutrient solution, either sucrose (Carl Roth GmbH & Co.) or yeast autolysate (MP Biomedicals). Phosphoric (0.01%) and propionic (0.1%) acids were added to both solutions as inhibitors of bacteria and mold growth. Tubes with flies were placed into closed container with moistened cotton tissue in the bottom to maintain high humidity and decrease evaporation of solutions from the capillaries. Tubes with capillaries but without the fly were used as the evaporation controls, and those values were subtracted from the experimental data. The amount of food consumed was tracked every 24 h during 6 consecutive days, and capillary tubes were replaced with new ones containing fresh solutions. From 14 to 25 flies were tested per genotype for each food treatment.

### Dietary Treatments

We used 9 dietary conditions, providing flies with one of three concentrations each (3, 6 and 12%) of yeast (the only source of protein offered, but also containing vitamins, minerals and carbohydrates) and sucrose solutions in separate capillaries. Rather than being mixed together, yeast and sucrose solutions were provided separately to allow flies to mix their intake to meet their particular physiological demands. This means that the flies were able to compensate for dilution of one macronutrient source (yeast or sugar), without having simultaneously to over-eat the other. The term “diet” in our study represents the concentration of nutrients provided in the test solutions, rather than actual intakes. For example, “diet” with 3% sucrose and 3% yeast or yeast autolysate (3S–3Y) does not necessarily mean that flies ingested equal amounts of yeast and sucrose. Similarly, “diet” 12S–12Y does not always imply consuming greater amounts of yeast and sucrose than on “diet” 3S–3Y. To lessen this confusion we mark a diet treatment as, e.g., 3S–3Y instead of 3S:3Y, which implies an ingested ratio.

When analyzing the impacts on metabolism of dietary treatments, we translated volumes ingested of the yeast and sucrose solutions into actual intakes of carbohydrate and protein, with carbohydrate coming both from yeast and sucrose. To calculate the amounts of carbohydrate and protein eaten, the volumes consumed were multiplied by the concentrations of solutions (30 mg/ml for 3%, 60 mg/ml for 6% and 120 mg/ml for 12% solutions). Carbohydrate content in yeast autolysate was 24% and protein, 45% (Lee et al., 2008).

### Metabolic Parameters

Flies were decapitated and then centrifuged (3000 g, 5 min) to extract hemolymph, which was used for determination of glucose and trehalose. Pre-weighed bodies were homogenized in 50 mM Nabuffer, centrifuged, and used for determination of glucose and glycogen level. Measurements were performed using a glucose assay kit (Liquick Cor-Glucose diagnostic kit, Cormay, Poland, Cat. № 2-203). The glycogen was converted into glucose by amylogucosidase from *Aspergillus niger* (25 °C, 4 hours). For triacylgyceride (TAG) determination, flies were weighted, homogenized in 20 mM PBST (phosphate buffered saline containing 0.05% Triton X-100), boiled and centrifuged (13000g, 10 min) (Rovenko et al., 2015a; Rovenko et al., 2015b). Resulting supernatants were used for the triglyceride assay with a Liquick Cor-TG diagnostic kit (Cormay, Poland, Cat. № 2-254). Flies of all genotypes were tested in 4-6 independent replicates (Rovenko et al., 2015a).

Total protein content in whole body was measured using Coomassie blue dye according to Bradford. Body supernatants were obtained by homogenization in 50 mM potassium phosphate buffer in ratio 1:10 and centrifuged (13000 g, 15 min, 4°C). Serum bovine albumin was a standard. Data are expressed as mg of protein per g of wet fly body weight (mg/gww).

### Data Visualization and Statistics

Statistical comparisons were performed using software GraphPad Prism 6 and additionally reassessed in R v.3.3.1. Analysis of variance (ANOVA) followed by Tukey’s test for multiple comparisons with Bonferroni correction was performed for all data. We compared values for either the different fruit fly lines subjected to one particular dietary treatment or between values for one particular line but fed on different dietary treatments. In the latter case, our ANOVA model included influence of dietary yeast or sucrose alone, or interaction between both. Boxplots represent median value (line inside box) with upper and lower quartiles (bottom and top borders of the box) for 75% and 25% of the data, respectively. Upper and lower whiskers represent 90% and 10% of the data, while values outside of the whiskers (outliers) are shown as dots.

Responses for metabolic variables were mapped onto bi-coordinate intakes of protein and carbohydrate, visualized using thin-plate splines and supported statistically using generalized additive models (GAMs) implemented in R (Solon-Biet et al., 2014).

## SUPPLEMENTAL DATA

Supplemental Information includes two figures and nine tables and can be found with this article online at

## AUTHOR CONTRIBUTIONS

Conceptualization, O.L., K.S., and S.S.; Methodology, O.L. and S.S.; Investigation, U.S., H.F., and I.Y.; Writing – Original Draft, D.G., O.L., U.S.; Writing – Reviews & Editing, O.L., S.S., K.S., A.V.; Formal Analysis, U.S., O.L., D.G.; Data curation, S.S.; Visualization, O.L., D.G., S.S.

## ACKNOWLEDGEMENTS

We thank Sebastian Grönke and Professor Dame Linda Partridge for providing us with lines of fruit flies used in this study; Samantha M. Solon-Biet and Rahul Gokarn for the help with calculation of generalized additive model estimates and response surface visualization; William Ja and J. Andrew Pospisilik for the critical reading of the manuscript.

## Supplemental data

**Figure S1.**
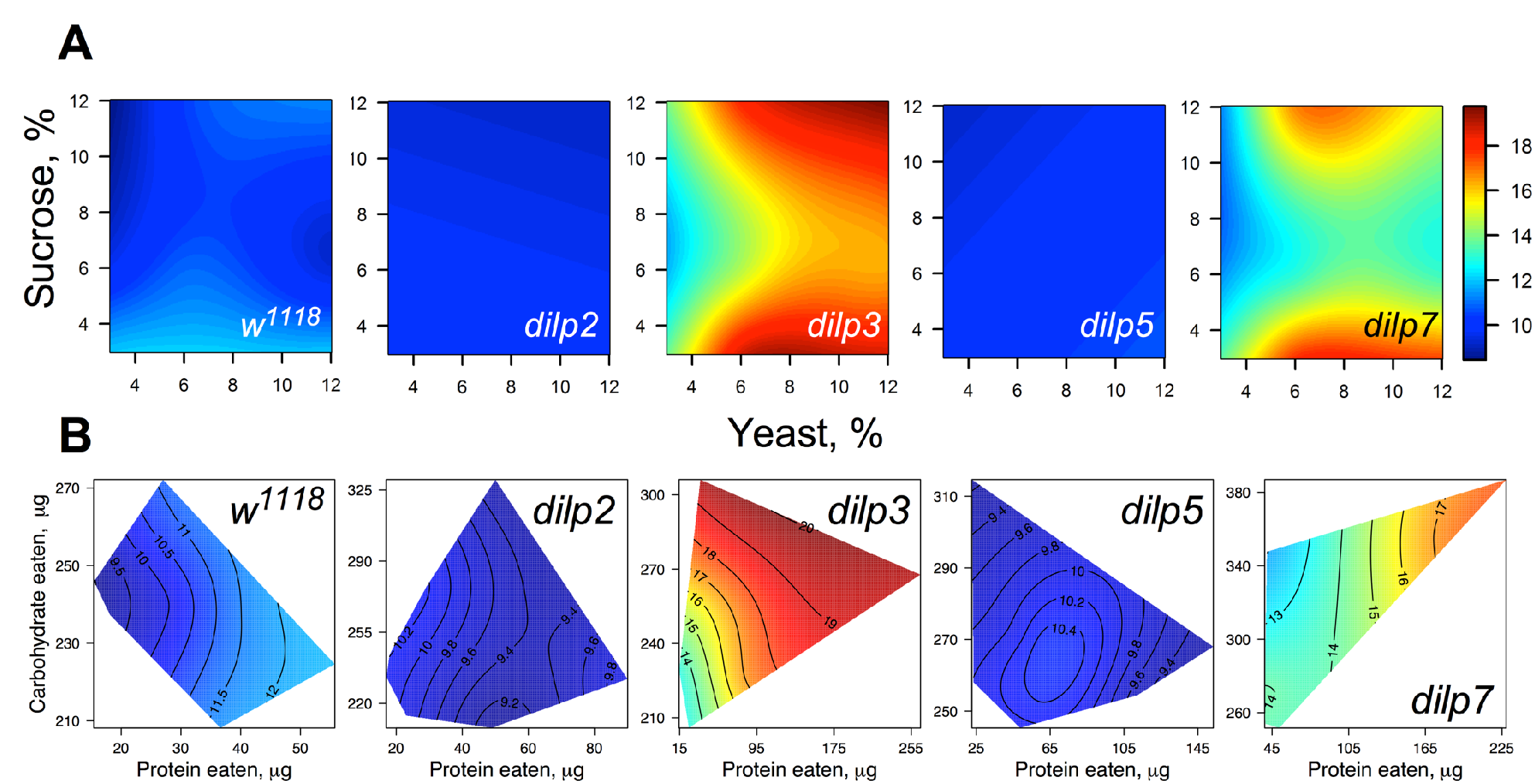
The diet-dependent patterns of body glucose in wild type and *dilp* mutant flies (A) Dietary response surfaces depicting dependence of glucose content in the body of individuals of wild type (*w^1118^*) and *dilp* mutant lines on concentrations of yeast and sucrose in the diet. (B) Dietary response surfaces showing dependence of body glucose content in wild type and *dilp* mutant lines on amounts of protein and carbohydrate consumed. Each surface has own scale shown by contour lines.

**Figure S2.**
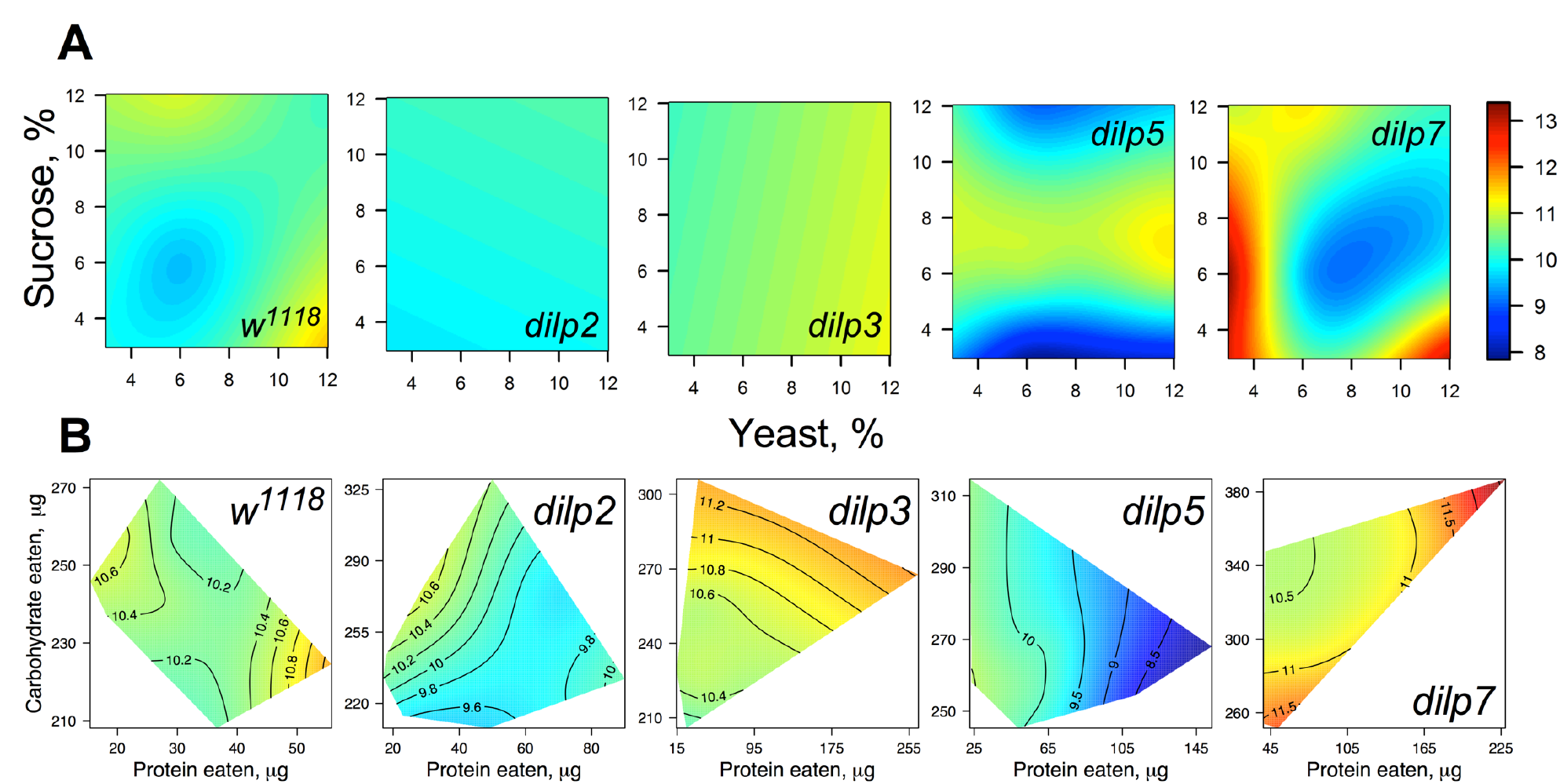
The diet-dependent patterns of body trehalose in wild type and *dilp* mutant flies (A) Dietary response surfaces depicting dependence of trehalose content in the body of individuals of wild type (*w^1118^*) and *dilp* mutant lines on concentrations of yeast and sucrose in the diet. (B) Dietary response surfaces showing dependence of body trehalose content in wild type and *dilp* mutant lines on amounts of protein and carbohydrate consumed. Each surface has own scale shown by contour lines.

**Table S1.** Coefficients for genotype-diet interactions provided by generalized linear models assessing contribution of genotype to volume of sucrose solution ingested or sucrose amount eaten.

|  | Sucrose volume |  |  |  | Sucrose amount |  |  |  |
| --- | --- | --- | --- | --- | --- | --- | --- | --- |
| | value | df | $\chi^2$ | p-value | value | df | $\chi^2$ | p-value |
| $w^{1118}$-dilp1 | 0.286 | 1.00 | 1.09 | 1.000 | 21.622 | 1.00 | 7.29 | 0.083 |
| $w^{1118}$-dilp2 | 0.005 | 1.00 | 0.00 | 1.000 | -12.640 | 1.00 | 2.76 | 0.775 |
| $w^{1118}$-dilp2,3 | -0.239 | 1.00 | 0.88 | 1.000 | -34.652 | 1.00 | 21.64 | 0.000 * |
| $w^{1118}$-dilp3 | -0.378 | 1.00 | 2.11 | 1.000 | -4.101 | 1.00 | 0.29 | 1.000 |
| $w^{1118}$-dilp4 | 0.194 | 1.00 | 0.41 | 1.000 | -27.167 | 1.00 | 9.39 | 0.033 * |
| $w^{1118}$-dilp5 | 0.090 | 1.00 | 0.12 | 1.000 | -25.253 | 1.00 | 11.44 | 0.012 * |
| $w^{1118}$-dilp7 | 0.752 | 1.00 | 7.54 | 0.157 | -60.351 | 1.00 | 56.77 | 0.000 * |
| dilp1-dilp2 | -0.281 | 1.00 | 0.84 | 1.000 | -34.262 | 1.00 | 14.58 | 0.003 * |
| dilp1-dilp2,3 | -0.525 | 1.00 | 3.02 | 1.000 | -56.274 | 1.00 | 40.58 | 0.000 * |
| dilp1-dilp3 | -0.092 | 1.00 | 0.09 | 1.000 | -25.723 | 1.00 | 8.22 | 0.054 |
| dilp1-dilp4 | 0.480 | 1.00 | 1.95 | 1.000 | -48.789 | 1.00 | 23.53 | 0.000 * |
| dilp1-dilp5 | 0.376 | 1.00 | 1.54 | 1.000 | -46.875 | 1.00 | 28.07 | 0.000 * |
| dilp1-dilp7 | 1.037 | 1.00 | 10.62 | 0.030 * | -81.973 | 1.00 | 77.50 | 0.000 * |
| dilp2-dilp2,3 | -0.244 | 1.00 | 0.71 | 1.000 | -22.013 | 1.00 | 6.75 | 0.103 |
| dilp2-dilp3 | -0.373 | 1.00 | 1.60 | 1.000 | 8.538 | 1.00 | 0.98 | 1.000 |
| dilp2-dilp4 | 0.199 | 1.00 | 0.36 | 1.000 | -14.527 | 1.00 | 2.22 | 0.952 |
| dilp2-dilp5 | 0.095 | 1.00 | 0.11 | 1.000 | -12.613 | 1.00 | 2.21 | 0.952 |
| dilp2-dilp7 | 0.757 | 1.00 | 6.08 | 0.342 | -47.711 | 1.00 | 28.28 | 0.000 * |
| dilp2,3-dilp3 | -0.617 | 1.00 | 4.53 | 0.764 | 30.551 | 1.00 | 12.99 | 0.006 * |
| dilp2,3-dilp4 | -0.045 | 1.00 | 0.02 | 1.000 | 7.486 | 1.00 | 0.61 | 1.000 |
| dilp2,3-dilp5 | -0.149 | 1.00 | 0.27 | 1.000 | 9.400 | 1.00 | 1.27 | 1.000 |
| dilp2,3-dilp7 | 0.513 | 1.00 | 2.88 | 1.000 | -25.699 | 1.00 | 8.46 | 0.051 |
| dilp3-dilp4 | -0.572 | 1.00 | 2.95 | 1.000 | -23.066 | 1.00 | 5.60 | 0.161 |
| dilp3-dilp5 | -0.468 | 1.00 | 2.60 | 1.000 | -21.151 | 1.00 | 6.21 | 0.127 |
| dilp3-dilp7 | -1.130 | 1.00 | 13.56 | 0.006 * | -56.250 | 1.00 | 39.30 | 0.000 * |
| dilp4-dilp5 | 0.104 | 1.00 | 0.10 | 1.000 | 1.914 | 1.00 | 0.04 | 1.000 |
| dilp4-dilp7 | -0.558 | 1.00 | 2.63 | 1.000 | -33.184 | 1.00 | 10.89 | 0.015 * |
| dilp5-dilp7 | -0.662 | 1.00 | 4.78 | 0.689 | -35.098 | 1.00 | 15.74 | 0.001 * |

**Table S2.** Coefficients for genotype-diet interactions provided by generalized linear models assessing contribution of genotype to volume of yeast autolysate solution ingested or yeast autolysate amount eaten.

|  | Yeast autolysate volume |  |  |  | Yeast autolysate amount |  |  |  |
| --- | --- | --- | --- | --- | --- | --- | --- | --- |
| | value | df | $\chi^2$ | *p*-value | value | df | $\chi^2$ | *p*-value |
| *w<sup>1118</sup>-dilp1* | -0.091 | 1.00 | 0.31 | 1.000 | -1.219 | 1.00 | 0.04 | 1.000 |
| *w<sup>1118</sup>-dilp2* | -0.121 | 1.00 | 0.60 | 1.000 | -10.526 | 1.00 | 3.60 | 0.404 |
| *w<sup>1118</sup>-dilp2,3* | -1.590 | 1.00 | 107.89 | 0.000 * | -50.307 | 1.00 | 85.96 | 0.000 * |
| *w<sup>1118</sup>-dilp3* | -0.776 | 1.00 | 24.57 | 0.000 * | -32.376 | 1.00 | 34.08 | 0.000 * |
| *w<sup>1118</sup>-dilp4* | -0.914 | 1.00 | 25.18 | 0.000 * | -25.669 | 1.00 | 15.80 | 0.001 * |
| *w<sup>1118</sup>-dilp5* | -1.137 | 1.00 | 54.89 | 0.000 * | -35.446 | 1.00 | 42.48 | 0.000 * |
| *w<sup>1118</sup>-dilp7* | -1.959 | 1.00 | 141.61 | 0.000 * | -55.870 | 1.00 | 91.70 | 0.000 * |
| *dilp1-dilp2* | -0.030 | 1.00 | 0.03 | 1.000 | -9.307 | 1.00 | 2.03 | 0.927 |
| *dilp1-dilp2,3* | -1.499 | 1.00 | 68.18 | 0.000 * | -49.088 | 1.00 | 58.20 | 0.000 * |
| *dilp1-dilp3* | -0.684 | 1.00 | 13.78 | 0.002 * | -31.158 | 1.00 | 22.73 | 0.000 * |
| *dilp1-dilp4* | -0.823 | 1.00 | 15.87 | 0.001 | -24.450 | 1.00 | 11.14 | 0.010 * |
| *dilp1-dilp5* | -1.046 | 1.00 | 33.07 | 0.000 * | -34.227 | 1.00 | 28.20 | 0.000 * |
| *dilp1-dilp7* | -1.868 | 1.00 | 95.27 | 0.000 * | -54.651 | 1.00 | 64.93 | 0.000 * |
| *dilp2-dilp2,3* | -1.469 | 1.00 | 71.16 | 0.000 * | -39.781 | 1.00 | 41.52 | 0.000 * |
| *dilp2-dilp3* | -0.655 | 1.00 | 13.66 | 0.002 * | -21.850 | 1.00 | 12.11 | 0.007 * |
| *dilp2-dilp4* | -0.794 | 1.00 | 15.71 | 0.001 * | -15.143 | 1.00 | 4.55 | 0.263 |
| *dilp2-dilp5* | -1.016 | 1.00 | 33.90 | 0.000 * | -24.919 | 1.00 | 16.23 | 0.001 * |
| *dilp2-dilp7* | -1.838 | 1.00 | 99.37 | 0.000 * | -45.344 | 1.00 | 48.13 | 0.000 * |
| *dilp2,3-dilp3* | 0.814 | 1.00 | 21.86 | 0.000 * | 17.930 | 1.00 | 8.43 | 0.037 * |
| *dilp2,3-dilp4* | 0.676 | 1.00 | 11.69 | 0.006 * | 24.638 | 1.00 | 12.37 | 0.006 * |
| *dilp2,3-dilp5* | 0.453 | 1.00 | 6.99 | 0.066 | 14.861 | 1.00 | 5.98 | 0.130 |
| *dilp2,3-dilp7* | -0.369 | 1.00 | 4.13 | 0.269 | -5.563 | 1.00 | 0.75 | 1.000 |
| *dilp3-dilp4* | -0.139 | 1.00 | 0.48 | 1.000 | 6.707 | 1.00 | 0.89 | 1.000 |
| *dilp3-dilp5* | -0.361 | 1.00 | 4.28 | 0.269 | -3.069 | 1.00 | 0.25 | 1.000 |
| *dilp3-dilp7* | -1.183 | 1.00 | 41.18 | 0.000 * | -23.493 | 1.00 | 12.92 | 0.005 * |
| *dilp4-dilp5* | -0.222 | 1.00 | 1.26 | 1.000 | -9.777 | 1.00 | 1.94 | 0.927 |
| *dilp4-dilp7* | -1.044 | 1.00 | 25.54 | 0.000 * | -30.201 | 1.00 | 16.99 | 0.001 * |
| *dilp5-dilp7* | -0.822 | 1.00 | 20.44 | 0.000 * | -20.424 | 1.00 | 10.04 | 0.017 * |

**Table S3.** Coefficients of the generalized additive models describing the response of circulating glucose in fruit flies of the lines investigated in this study to concentrations of sucrose and yeast in proposed food.

| | $w^{1118}$ | *dilp2* | *dilp3* | *dilp5* | *dilp7* |
| --- | --- | --- | --- | --- | --- |
| edf |  |  |  |  |  |
| S | 0.7843 | 1.3690 | 0.0001 | 0.7418 | 0.0001 |
| Y | 0.0001 | 0.5779 | 1.5576 | 0.6503 | 1.3210 |
| S × Y | 0.7292 | 0.0000 | 0.4737 | 0.0001 | 1.4120 |
| ref. df |  |  |  |  |  |
| S | 2.0 | 2.0 | 2.0 | 2.0 | 2.0 |
| Y | 2.0 | 2.0 | 2.0 | 2.0 | 2.0 |
| S × Y | 3.0 | 3.0 | 3.0 | 3.0 | 3.0 |
| F |  |  |  |  |  |
| S | 1.817 | 1.969 | 0.000 | 1.436 | 0.000 |
| Y | 0.000 | 0.685 | 6.006 | 0.930 | 2.830 |
| S × Y | 0.507 | 0.000 | 0.214 | 0.000 | 1.331 |
| p-value |  |  |  |  |  |
| S | 0.0261 | 0.0711 | 0.4778 | 0.0572 | 0.7685 |
| Y | 0.6007* | 0.1324 | 0.0011* | 0.1000 | 0.0117* |
| S × Y | 0.1299 | 0.5411 | 0.2474 | 0.5014 | 0.0528 |

Here and in the tables below:
edf – estimated degrees of freedom
Ref. df – degrees of freedom for reference distribution

**Table S4.**
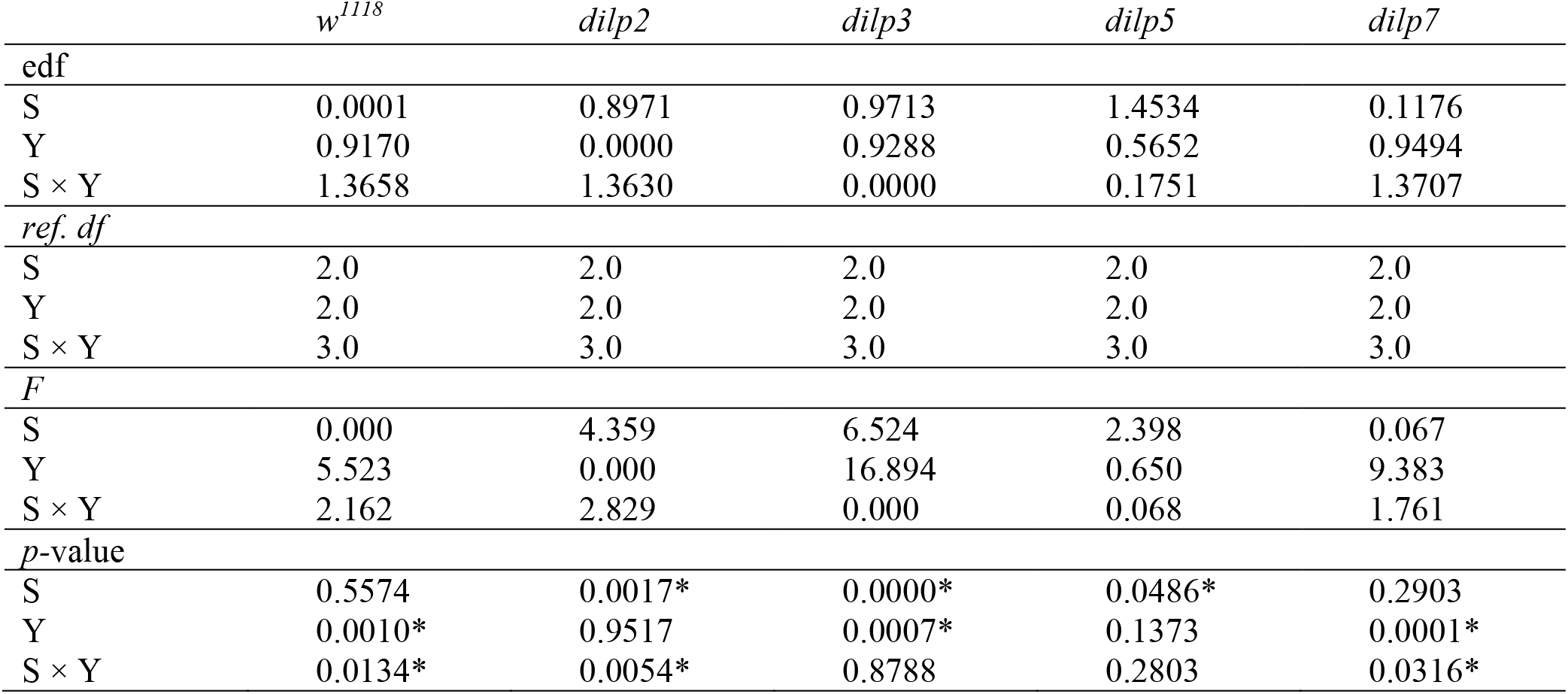
Coefficients of the generalized additive models describing the response of circulating trehalose in fruit flies of the lines investigated in this study to concentrations of sucrose and yeast in proposed food.

**Table S5.** Coefficients of the generalized additive models describing the response of glycogen content in fruit flies of the lines investigated in this study to concentrations of sucrose and yeast in proposed food.

| | $w^{1118}$ | *dilp2* | *dilp3* | *dilp5* | *dilp7* |
| --- | --- | --- | --- | --- | --- |
| edf |  |  |  |  |  |
| S | 1.5600 | 0.0000 | 1.6260 | 0.0001 | 0.9041 |
| Y | 0.0000 | 0.0000 | 0.5468 | 1.3000 | 0.0001 |
| S × Y | 0.0000 | 0.0000 | 0.0000 | 1.2170 | 1.6031 |
| Ref. df |  |  |  |  |  |
| S | 2.0 | 2.0 | 2.0 | 2.0 | 2.0 |
| Y | 2.0 | 2.0 | 2.0 | 2.0 | 2.0 |
| S × Y | 3.0 | 3.0 | 3.0 | 3.0 | 3.0 |
| *F* |  |  |  |  |  |
| S | 5.629 | 0.000 | 12.159 | 0.000 | 4.714 |
| Y | 0.000 | 0.000 | 0.603 | 2.593 | 0.000 |
| S × Y | 0.000 | 0.000 | 0.000 | 1.538 | 3.423 |
| *p*-value |  |  |  |  |  |
| S | 0.0018* | 1.0000 | 0.0000* | 0.5807 | 0.0000* |
| Y | 0.9731 | 1.0000 | 0.1470 | 0.0270* | 0.4413 |
| S × Y | 0.9995 | 1.0000 | 0.4620 | 0.0308* | 0.0036* |

**Table S6.** Coefficients of the generalized additive models describing the response of triacylglyceride content in fruit flies of the lines investigated in this study to concentrations of sucrose and yeast in proposed food.

| | $w^{1118}$ | *dilp2* | *dilp3* | *dilp5* | *dilp7* |
| --- | --- | --- | --- | --- | --- |
| edf |  |  |  |  |  |
| S | 0.9335 | 0.9445 | 0.0000 | 0.0000 | 0.8101 |
| Y | 0.0000 | 0.8561 | 0.4947 | 0.5962 | 0.8291 |
| S × Y | 0.2901 | 0.0001 | 0.0000 | 0.0003 | 2.1608 |
| Ref. df |  |  |  |  |  |
| S | 2.0 | 2.0 | 2.0 | 2.0 | 2.0 |
| Y | 2.0 | 2.0 | 2.0 | 2.0 | 2.0 |
| S × Y | 3.0 | 3.0 | 3.0 | 3.0 | 3.0 |
| *F* |  |  |  |  |  |
| S | 7.016 | 8.507 | 0.000 | 0.000 | 2.133 |
| Y | 0.000 | 2.973 | 0.489 | 0.738 | 2.424 |
| S × Y | 0.122 | 0.000 | 0.000 | 0.000 | 2.811 |
| *p*-value |  |  |  |  |  |
| S | 0.0003* | 0.0001* | 0.8060 | 1.0000 | 0.0260* |
| Y | 0.9351 | 0.0123* | 0.1680 | 0.1250 | 0.0194* |
| S × Y | 0.2753 | 0.5186 | 0.8980 | 0.3560 | 0.0168* |

**Table S7.**
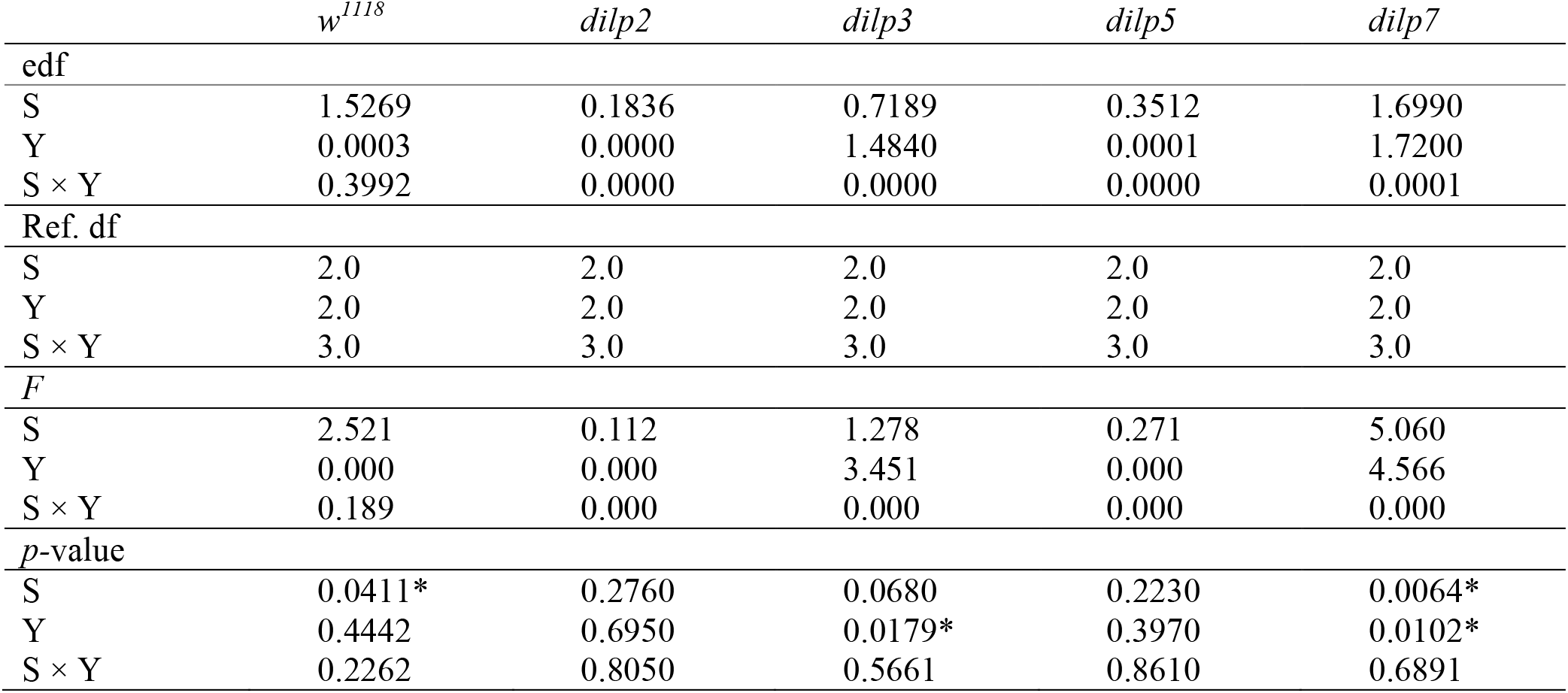
Coefficients of the generalized additive models describing the response of whole body glucose content in fruit flies of the lines investigated in this study to concentrations of sucrose and yeast in proposed food.

**Table S8.** Coefficients of the generalized additive models describing the response of whole body trehalose content in fruit flies of the lines investigated in this study to concentrations of sucrose and yeast in proposed food.

| | $w^{1118}$ | *dilp2* | *dilp3* | *dilp5* | *dilp7* |
| --- | --- | --- | --- | --- | --- |
| edf |  |  |  |  |  |
| S | 0.0000 | 0.0000 | 0.0000 | 0.5899 | 0.0011 |
| Y | 0.0000 | 0.0000 | 0.0001 | 0.0006 | 0.0002 |
| S × Y | 0.7184 | 0.0001 | 0.0000 | 0.0000 | 0.0001 |
| Ref. df |  |  |  |  |  |
| S | 2.0 | 2.0 | 2.0 | 2.0 | 2.0 |
| Y | 2.0 | 2.0 | 2.0 | 2.0 | 2.0 |
| S × Y | 3.0 | 3.0 | 3.0 | 3.0 | 3.0 |
| *F* |  |  |  |  |  |
| S | 0.000 | 0.000 | 0.000 | 0.719 | 0.001 |
| Y | 0.000 | 0.000 | 0.000 | 0.000 | 0.000 |
| S × Y | 0.458 | 0.000 | 0.000 | 0.000 | 0.000 |
| *p*-value |  |  |  |  |  |
| S | 0.7580 | 0.8090 | 0.9200 | 0.1280 | 0.3300 |
| Y | 0.6210 | 0.9720 | 0.3940 | 0.3340 | 0.4080 |
| S × Y | 0.1580 | 0.4300 | 0.4300 | 0.6320 | 0.7920 |

**Table S9.** Coefficients of the generalized additive models describing the response of weight of fruit flies of the lines investigated in this study to concentrations of sucrose and yeast in proposed food.

| | $w^{1118}$ | *dilp2* | *dilp3* | *dilp5* | *dilp7* |
| --- | --- | --- | --- | --- | --- |
| edf |  |  |  |  |  |
| S | 0.8210 | 0.8143 | 0.0000 | 0.0000 | 0.0000 |
| Y | 0.7545 | 0.0000 | 0.5382 | 0.0000 | 0.0000 |
| S × Y | 0.0000 | 0.0000 | 0.0000 | 0.0000 | 0.6730 |
| Ref. df |  |  |  |  |  |
| S | 2.0 | 2.0 | 2.0 | 2.0 | 2.0 |
| Y | 2.0 | 2.0 | 2.0 | 2.0 | 2.0 |
| S × Y | 3.0 | 3.0 | 3.0 | 3.0 | 3.0 |
| *F* |  |  |  |  |  |
| S | 2.294 | 1.414 | 0.583 | 0.000 | 0.000 |
| Y | 1.536 | 0.000 | 0.000 | 0.000 | 0.000 |
| S × Y | 0.000 | 0.000 | 0.000 | 0.000 | 0.381 |
| *p*-value |  |  |  |  |  |
| S | 0.0218* | 0.0610 | 0.1500 | 0.7940 | 0.4670 |
| Y | 0.0487* | 0.9750 | 0.9490 | 0.5910 | 0.8710 |
| S × Y | 0.9404 | 0.6710 | 0.6790 | 1.0000 | 0.1940 |

